# Multi-class Cancer Classification and Biomarker Identification using Deep Learning

**DOI:** 10.1101/2020.12.24.424317

**Authors:** Fariha Muazzam

**Affiliations:** Department of Computer Science, National University of Computer And Emerging Sciences

**Keywords:** Cancer Detection, Cancer Prevention, Targeted Therapy, Precision Medicine

## Abstract

Genetic data is important for analysing cellular functions whose disruption gives rise to various kinds of cancer. The intricacies of gene interaction are captured in various kinds of data for cancer detection through sequencing technology, but diagnosis, prognosis and treatment are still hard. Advent of machine learning helped researchers in supervised and unsupervised learning tasks along with gene identification but resourcefulness has not been overtly satisfactory. This research revolves around multi-class cancer classification, feature extraction and relevant gene identification through deep learning methods for 12 different types of cancers using RNA-SEQ from The Cancer Genome Atlas.

It has been constrained by hardware resource availability and within them the experiments that have been performed have shown promising results. Stacked De-noising Autoencoders were used for feature extraction and biomarker identification while 1D Convolutional Neural Networks for classification. Classification was performed with extracted features and relevant genes, which gave average performance of around 94% and 95% respectively. We were able to identify generic cancer-related pathways and their associated genes through Stacked De-noising Auto-encoders generated weight matrix and features. The common pathways include WNT Signalling Pathway, Angiogenesis. Moreover, across all pathways some recurrent genes were observed, namely: PIK3C2G, PCDHB8, WNT10A and these genes were found, in literature, to be involved in multiple types of cancer.

The proposed approach shows superior performance and promise against traditional techniques used by bioinformatics community, in terms of accuracy and relevant gene identification.

## INTRODUCTION

Genes play an important role in the normal functioning of humans’ bodily processes and physiology (1). However, there is a nuance of uncertainty associated with molecular events that occur which can cause alteration in routine processes. Such changes in mechanism can lead to mutations or chromosomal rearrangements which can be harmful or benign, but are heavily associated with cancer causation (1). Identification of genes or group of genes propagating cancerous cell formation provides meaningful opportunity to detect cancer at an early stage or stagnate its progression at a later stage (1).

In today’s day and age cancer is one of the leading diseases, causing 8.2 million deaths each year (2). Cancer diagnosis and treatment remain to be center of attention for medical professionals and researchers everywhere. Development of high-throughput DNA sequencing technology has led to varied discoveries in the field of genomics as mutation profiles, RNA expressions or micro-RNA profiles can be easily detected now (1).The importance of such genetic data can be realized by the fact that cancer diagnosis, progression and prognosis can be statistically analyzed through machine learning algorithms. Furthermore, sub-networks of genes and individual biomarkers responsible for cancer can be marginalized for precision medicine (1) (3).

Machine learning and deep learning techniques have been used extensively in domains such as image processing, natural language processing or audio recognition and have shown great promise. However, with regard to field of bioinformatics, focus has always been towards recognizing subtypes or biomarkers through clustering algorithms. In recent past, focus has shifted towards classification through supervised learning algorithms for RNA-seq expressions. With somatic mutations, very naive or basic methods have been used for classification. Also, multi-class classification has not really been explored even though cross-cancer biomarkers identification has been tampered with.

Machine learning algorithms ease two challenges associated with study of genetic data: extraction of meaningful genes and classification of cancer. Techniques like Principal Component Analysis, K-Means Clustering and Independent Component Analysis have been used to reduce dimensions while K-Nearest Neighbors, Random Forrest and Support Vector Machines for classification (4) (5). Due to availability of large datasets and computational resources, researchers have moved towards using deep learning algorithms in classification problems like object detection or image classification (1). More recently, bioinformatics has been penetrated with the applications of deep learning to genetic data for drug discovery, gene regulation or protein classification as huge sets of data are accessible (6). Hence, cancer detection based on gene expressions or mutation profiles has been experimented with deep learning architectures to improve classification accuracy and identification of biomarkers.

For cancer detection through gene expressions, Generative Adversarial Network(GAN) (7), Stacked Denoising Auto encoder(SDA)(5), Artificial Neural Networks(ANN) (5), Discriminant Deep Belief Networks(DDBN)(8) and One-Dimensional Convolutional Neural Networks(1DCNN) (9) have been used. Deep Neural Networks(DNN) have been used for heterogeneous classification of different types of cancer using somatic mutation profiles (1).

This research picks up from detection of different types of cancer RNA-Seq expressions using deep neural networks with application of dimensionality reduction (1). RNA-seq expressions data for breast cancer has been reduced using Kernel Principal Component Analysis(KPCA) and Principal Component Analysis(PCA) and classified using SVM with Linear and Radial Basis Function kernels and ANN (5). Heterogeneous RNA-seq expressions data has been analyzed with SDA for feature extraction and biomarker identification and DNN and 1DCNN for multi-class classification. The purpose was to achieve high classification performance and extract meaningful genes for targeted therapy by exploring deep learning architectures that have not been tried yet on RNA-seq expressions based cancer classification.

This section would be followed by description of materials and methods, results acquired and final conclusion of the whole study.

## RELATED WORK

Cancer detection from genetic data has been a challenging task but an important one for bioinformatics researchers. Due to cheaper DNA sequencing technology, larger datasets are available to be used for diagnosis, treatment or prognosis. Hence, various feature extraction and machine learning algorithms have been used for dimensionality reduction and classification over the years. Moreover, the world has moved from studying effects of individual gene functions to gene networks. Moreover, same networks can cause various diseases as well.

Over the years with advancement of sequencing technology, scientists have incorporated various forms of gene expressions data in their studies; ranging from microarray expression to DNA sequencing (10). In recent years the shift has been moved from microarrays to RNA-seq datasets for gene expressions-based cancer research. However, regardless of the data type most of the techniques used for cancer detection or relevant gene identification have been the same.

Clustering analysis has been used to group significant genes together and aid with accurate classification of samples. K-Nearest Neighbours has been used for quantifying correlation between gene expressions for prostate cancer (11) and with varied distance measures for classification of breast cancer (12). Also k-means clustering classification based on driver genes, identified using wavelet transforms for colon and leukemia samples (13). Hierarchal clustering has been utilized to classify subtypes of breast cancer data(14) and cancer data with reduced dimensionality(15). Apart from clustering, SVMs have been used stupendously for classification of gene expression profiles for different kinds of cancers. Multi-category SVMs aided the subtype classification of leukaemia dataset to a great extent (4). Network Algorithms have also been used to identify network of genes contributing to propagation of multiple types of cancer (16).

However, since, numerous machine learning algorithms have been developed; researchers have explored their usefulness with respect to cancer diagnosis and biomarker identification. With the advent of deep learning methods, there has been an obvious inclination towards using them for dimensionality reduction as well as classification.

Gupta et al.(1) in their paper used this architecture for learning meaningful representation of gene expressions data of yeast cell cycle Clusters of genes evaluated from raw input were already labeled and were compared with the clustering of output of SDA. Moreover, PCA was also tested on gene expression profiles and evaluated with aforementioned clustering algorithms. The results reveal that SDA capture gene co-expressions better than PCA by all means.

Danaae et al. (5) focused their research on extracting deeply connected genes from RNA-seq expressions of breast cancer data using SDA. PCA and KPCA were used as comparative techniques to measure SDA’s efficacy. Apart from reducing dimensionality, they have analyzed the weight matrix of SDA to identify contributor genes. These genes have been tagged as Deeply Connected Genes(DCGs). Panther pathways was used to analyze functions corresponding to different genes and tumor suppressor genes.

Bhat et al. (7) experimented with Generative Adversarial Deep Convolution Networks to accurately classify gene expression-based datasets of two types of cancer: breast cancer and prostrate cancer.

Karabulut et al. (8) demonstrated the efficiency of DDBN on classification of cancer as compared to traditional model like SVM. Experiments were performed on three different types of cancer individually: laryngeal, colorectal and bladder. For comparison SVM, Random Forrest and K-NN were applied on all datasets. Results revealed that DDBN outperformed all the afore-mentioned classification.

Liu et al. (9) focused their research on discrimination of tumor samples from normal ones. They proposed sample expansion method inspired from SAE and SDA to enlarge training samples. 1DCNN has been proposed in this paper for tumor classification. It takes input in one dimensional vector instead of traditional two dimensions used for image classification. The performance of 1DCNN was better than that of SAE on each dataset

Teixeira et al. (17) worked for singling out most informative genes using SDA for classification of thyroid cancer using ANN. They used traditional methods like PCA and Kernel PCA for comparison with deep learning method for feature extraction. Output of SDA was analyzed by extracting the weight matrix and using Connected Weights Method and three groups of genes were discovered with inter-related functions.

Hence, the effectiveness of deep learning models for feature extraction and relevant gene identification is prominent especially when the world is moving towards precision medicine. So it is that, multi-class classification and biomarker identification is the current focus and for that reason researchers have been experimenting with deep learning. Deep learning has become famous for classification problems related to larger datasets and feature extraction for wide variety of fields and more recently for bioinformatics too.

## MATERIAL AND METHOD

### Acquisition of data

Gene Expressions datasets have been most widely used with relation to anomaly classification as mentioned in before. The dataset for this study has been formulated from The Cancer Genome Atlas(TCGA) supported portals.

#### RNA-seq Expressions

TCGA portal provides gene expressions data in form of read counts as well as normalized expressions for 33 different types of cancer. For multi-class classification, each kind of data has to have same genes and this is ensured by the fact that they are sequenced by same technology and preprocessed with same techniques. Broad Institute GDAC portal provides dataset for RNA-seq expressions in raw form as well as RSEM normalized form. For this research, Illumina Hiseq RSEM normalized dataset has been used as seen in Table 1

**Table 1:** TCGA multi-class cancer dataset

| Cancer Type | No. of Cancerous Samples |
| --- | --- |
| Breast invasive carcinoma(BRCA) | 1100 |
| Adrenocortical carcinoma(ACC) | 70 |
| Cervical and endocervical cancer(CESC) | 304 |
| Head and neck Squamous Carcinoma(HNSC) | 520 |
| Kidney renal papillary cell carcinoma(KIRP) | 290 |
| BrainLower Grade Glioma ( LGG) | 516 |
| Lung adenocarcinoma (LUAD) | 501 |
| Pancreatic adenocarcinoma (PAAD) | 178 |
| Prostate adenocarcinoma (PRAD) | 497 |
| Stomach adenocarcinoma(STAD) | 416 |
| Uterine Carcinosarcoma (UCS) | 57 |
| Bladder urothelial carcinoma(BLCA) | 408 |

### Dataset Split

The dataset for 12 types was combined into 1 dataset with each sample given a corresponding label for its type of cancer. The labels were numbers between 0-11 for each sample, where each number corresponds to a specific cancer type. The dataset contained around 4967 samples for 12 types of cancer, and was split into training, validation and test sets. The percentage split of each set was 70%, 15% and 15% respectively. As there was an apparent class imbalance among different types, so division of dataset was kept proportionate per class. To elaborate it means, that each type was divided into three sets with the afore-mentioned percentage split.

### Preprocessing

The genes have been normalized and those with zero values across all samples have been removed, as they would not contribute to the results.

### SDA

The experiments used output of SDA as an input to 1DCNN for classification of cancer types. SDA has been trained through greedy-layer wise training where each layer is trained for a specific number of iterations and the output of the preceding was used as input to the succeeding layer. Number of hidden units per layer were decreased gradually because it has known to incorporate the features better. Five experiments were performed revealing substantial results and they produced five high-ranked gene sets and reduced feature sets. The output of SDA was used in two ways:

#### Using Reduced Features

The output of the final layer of SDA was the reduced features of the dataset. These features for each sample were stored for training dataset after the desired iterations were performed. Final weights for each layer were also stored so that it could be used to reduce features for test and validation dataset.

#### Using High Ranked Genes

Final weight matrix when analyzed shows that the weights of the genes were normally distributed. A small portion of genes had high weights which had been regarded as high-weight genes. These genes were filtered in training and testing datasets so as to reduce the number of features as done in (18).

The weight matrix of each layer was multiplied to generate a Number of Genes X Number of Features Matrix.

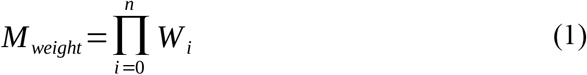

For each node, mean weight and standard deviation was calculated and genes were ranked by filtering genes outside specific number of standard deviations.

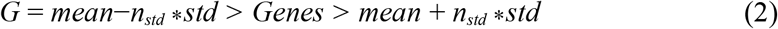

### 1DCNN

The reduced features extracted using SDA was fed to 1dcnn for classification. The overall accuracy of the system determines whether the extracted features were of any significance or not.

### Biomarker Identification

For biomarker identification, high-ranked gene sets were generated for different SDA architectures and their relevant pathways were identified from panther database. Overlapping pathways and genes were analyzed amongst all sets and there were quite a few that overlapped. The overlapping genes were checked against literature to confirm whether the identified genes are cross-cancer ones, and they are identified as biomarkers.

## RESULTS

This study focused exploration with RNA-Seq expression dataset only due to its availability; however it can be safely assumed that the built pipeline could be useful for other types of datasets as well.

### SDA

As mentioned in previous section, the output as reduced features and high-ranked genes based on weight matrix was used. Different number of layers of SDA was trained with different hidden units. As per literature, if the number of hidden units is decreased gradually then SDA better incorporates the features for reconstruction. The original number of genes was 20531 and removing the genes with zero value across all samples, left total number of genes to be 20313. The hidden units ranged between 15000-200 for whole architecture but first two layers contained fixed number of units 15000 and 10000 respectively. Only third-last and last layer were changed for experiments. Substantial experiments were conducted with 3 and 4 layers as that gave higher accuracy.

For reduced features, the best results were obtained when the reconstruction layer contained higher number of units. The features were tested by using 1DCNNs for classification. However, the accuracy kind of plateaued at 4000 features with around 96.5%.

The following graph in Fig 1 shows the accuracy achieved with 1dcnns and varied number of layers and reduced features. The experiments included in this graph are with 3 and 4 layers. The first and second layer contained fixed 15000 and 10000 units respectively.

**Figure 1:**
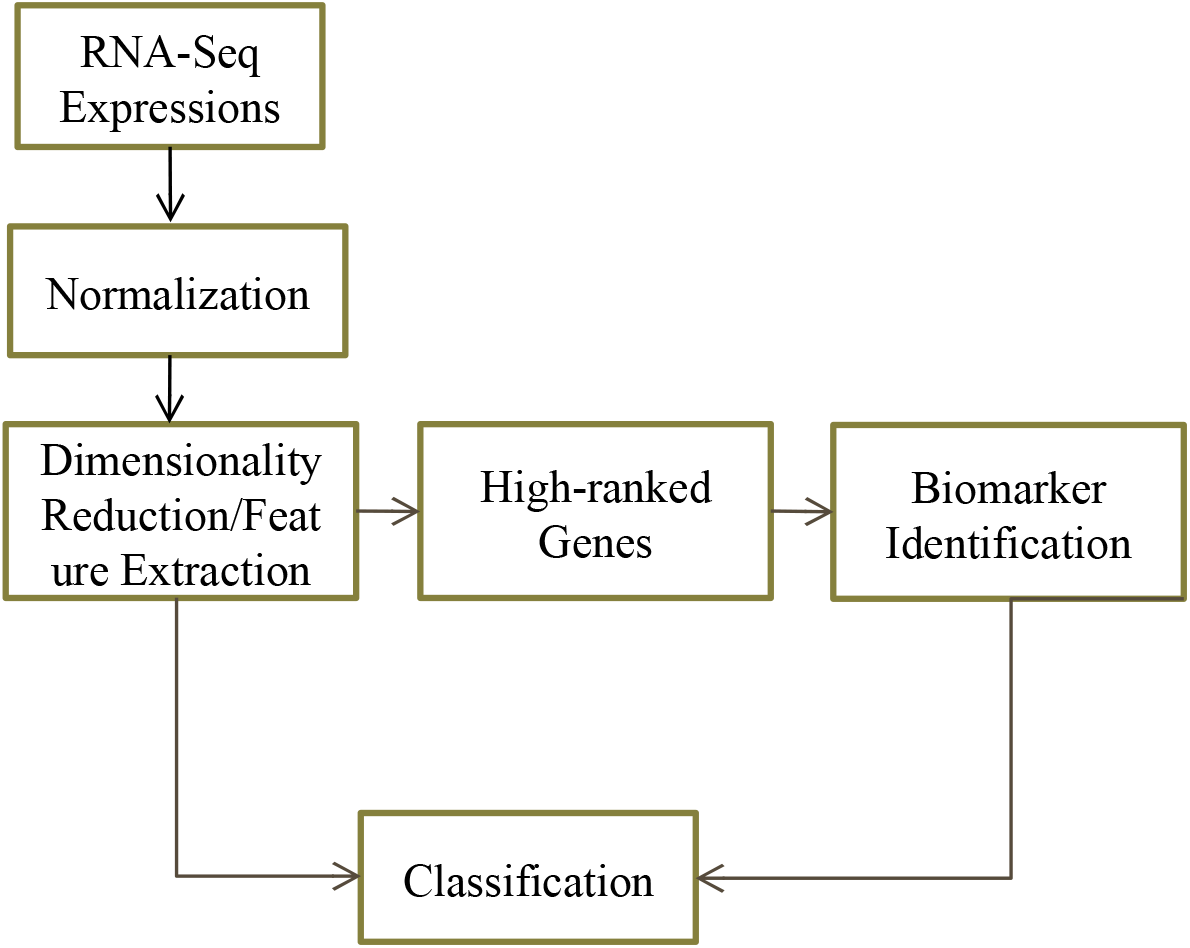
Workflow

### High-Ranked Genes

The weight matrix for each layer of SDA was used to rank genes based on the combination of their weights. As per literature it has been observed that genes with higher weights tend to act as contributing genes towards cancer. As per (18) the weight matrix of SDA follows an approximate normal distribution and the highly negative or highly positive genes in terms of their weights are significant genes. So, the genes away from mean weights would be categorized as the high-ranked ones. So we used standard deviation from the mean to identify the relevant genes. Due to limitation of resources, the experiments could only be performed within a restricted range; nevertheless they show huge performance in terms of relevant gene identification.

It was observed that genes that stood ground away from the mean were actually the relevant ones. Also the genes that overlapped amongst different SDA architectures were considered to be cross-cancer relevant genes. Since the aim of this research has always been that we achieve maximum performance with minimal genes; architectures within the range of 200-1000 features give better performance within 4-5 standard deviation.

Four genes were found to be similar amongst all pathways for all sets across all standard deviations, so proof of them being involved in multiple types was studied in literature. The study shows the promise and relevance of realized genes as seen in Table 2.

**Table 2:** Relevance of Identified Genes In Literature

| Genes | Cancer Types |
| --- | --- |
| WNT10A | BRCA <sup>19</sup><br>LUAD <sup>20</sup><br>BLCA <sup>21</sup><br>PRAD <sup>22</sup><br>PAAD <sup>23</sup> |
| PIK3C2G | BRCA <sup>24</sup><br>BLCA <sup>25</sup><br>HNSC <sup>26</sup> |

**Table 3:** Summarized Results for Reduced Features

| Paper | Classification | Mean Per Class Accuracy |
| --- | --- | --- |
| Danae. et al<sup>(5)</sup> | Breast Cancer | 98.26 |
| Proposed | Multi-class( 12 types including breast cancer) | 94.25 |

**Table 4:**
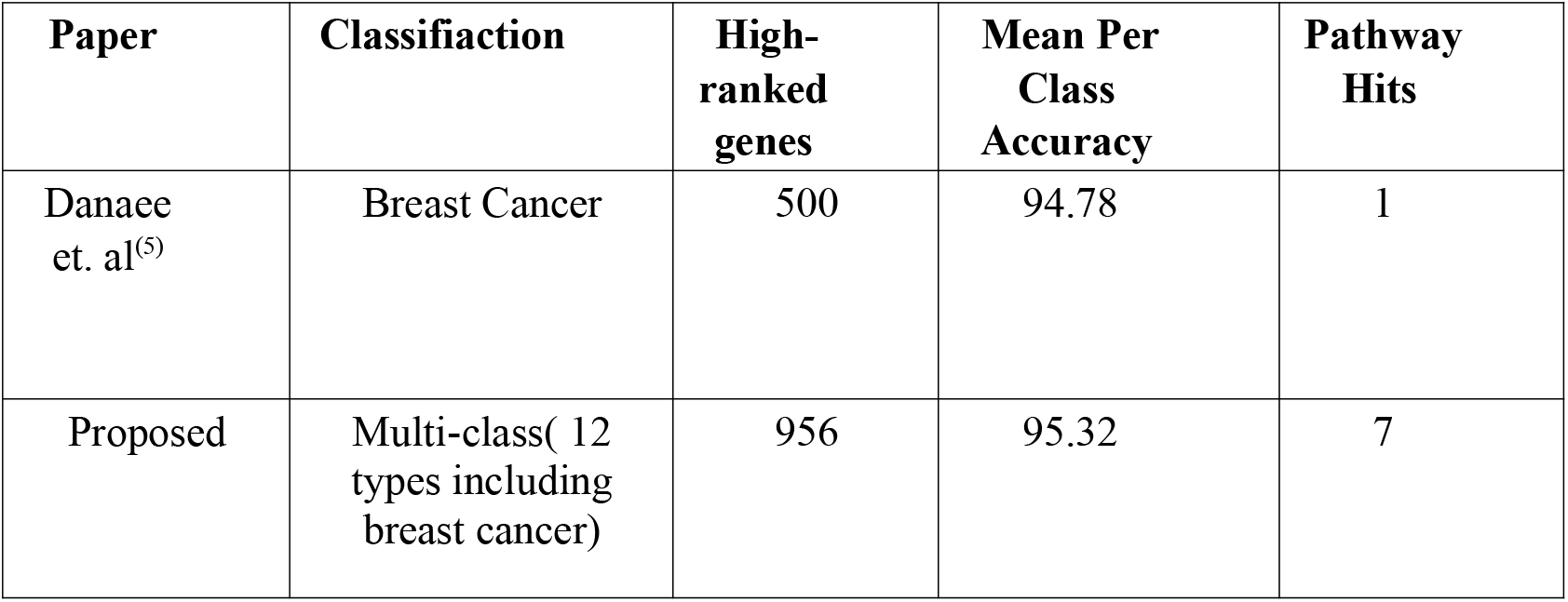
Summarized Results for High-ranked genes

Apart from that, there are 4 pathways that are found to be common in overlapping genes for different standard deviations, however two of them are same as found in all sets of experiment-generated genes for all standard deviations namely: WNT Pathway and Angiogenesis. Also the genes associated with these pathways are similar to that found in experiment-generated gene sets. The following Figure 3 shows how standard deviations between 4-5 relates pathways and overlapping genes and the scope for meaningful analysis.

**Figure 2:**
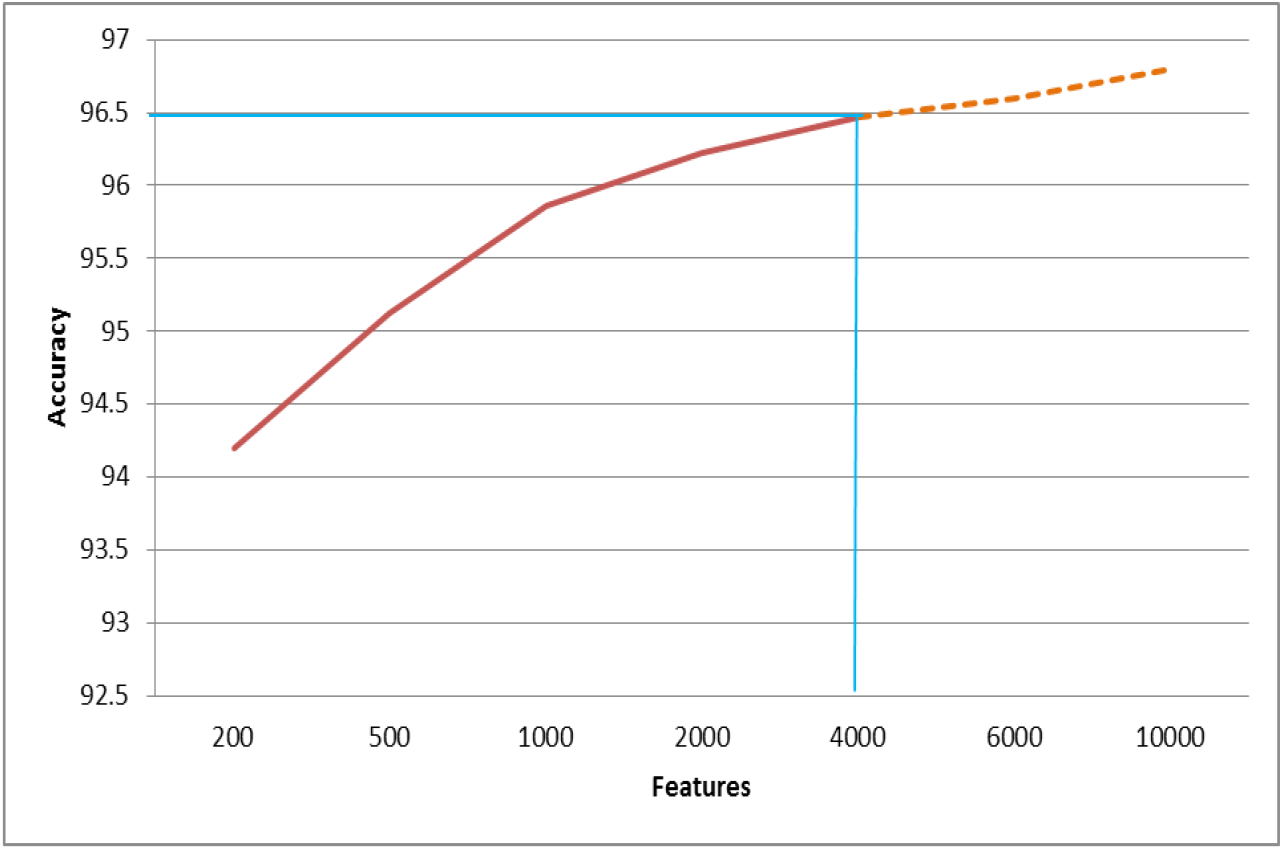
Accuracy with linear combination of reduced features

**Figure 3:**
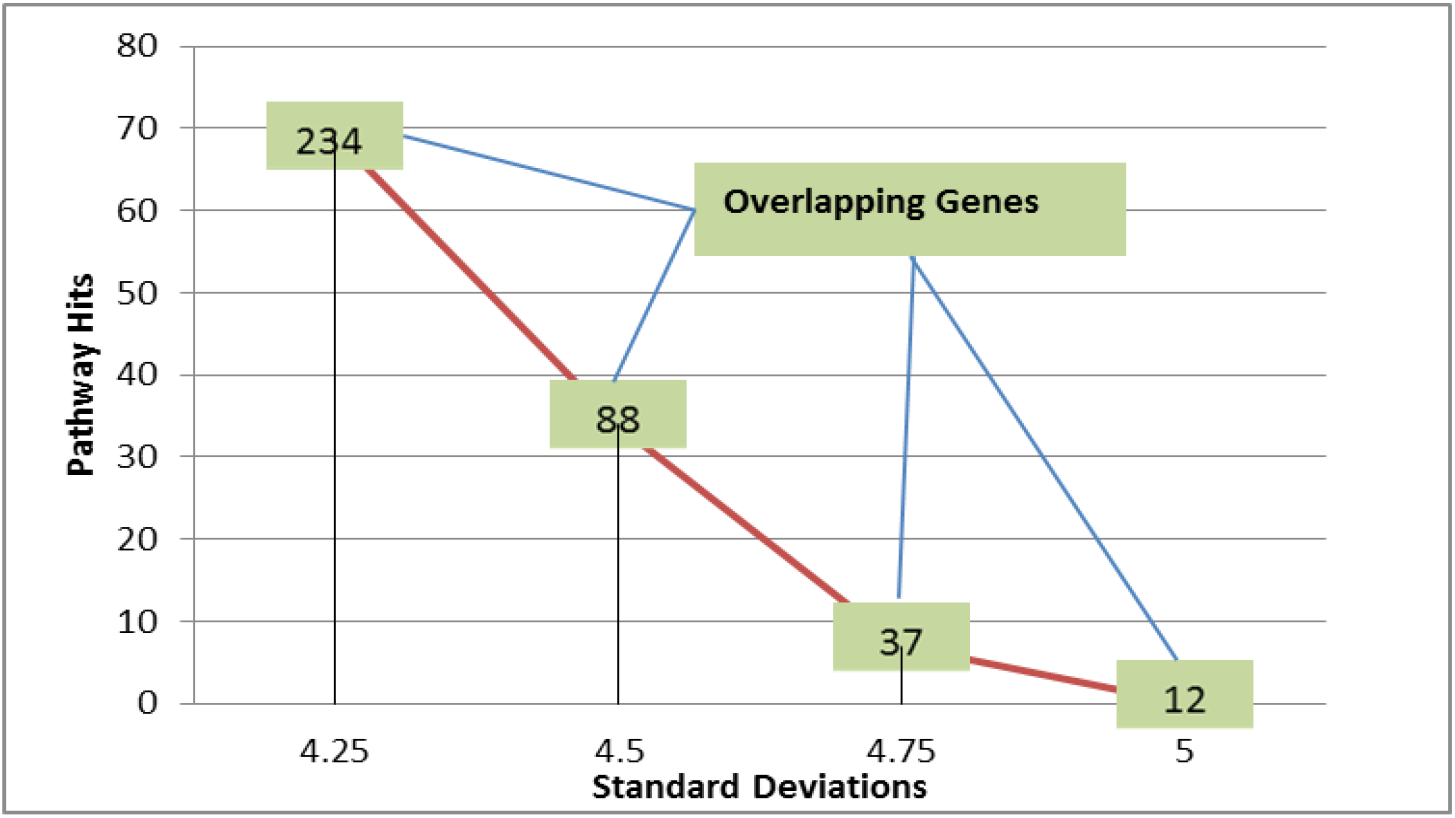
Pathway hits against different standard deviations

The following tables show the summarized results for reduced features and high-ranked genes in comparison to other similar studies.

## CONCLUSION

This study was aimed at classifying 12 types of cancer and identifying relevant genes and the results show that the proposed approach shows promise for the said task. Usage of SDA with 1DCNN has revealed an average accuracy of 94% for reduced features and 95% for high-ranked genes. This shows that relevant gene sets could help with cancer classification task as well as cross-cancer gene and pathway identification. We were able to identify cancer-relevant pathways and genes for the sets, that different experiments generated, from Panther Database. The common genes amongst all experiments were verified by literature as to be involved in multiple cancers. This shows that our method can be used for multi-class or single-class cancer classification and for recognizing the relevant genes as biomarkers. This gives hope to identify those genes that have yet not been explored by literature.

Panther Database is used by bioinformatics community to study the origin, families and relevance of genes with respect to single type or varied types of cancer. That involves a lot of manual analysis, but deep learning decreases the load by pointing to relevant genes and pathways or identify newer pathways and genes.

The hardware resource constrained the study but reliability and significance of automating the classification and identification with deep learning was still realized. More experiments would show more avenues that could be explored for cancer study through deep learning. Furthermore, using more types of cancer would also aid in identifying larger sets of cross-cancer biomarkers and pathways.

This study is just a step to show the relevance of using automated gene identification techniques which are reliable and can handle large amount of variations and unknowns and ambiguities. Whereas, the traditional statistical techniques for genes involve thresholding depending on the samples and the genes involved. Even though resource limitation in terms of GPU hours was tackled during the course of study, it still provided good results.

## ADDITIONAL INFORMATION

### Ethics Approval

This is an original study performed using open source dataset of TCGA and there is no violation of rights and obligations for usage of the dataset.

### Data Availability

The data was downloaded from broad institute firehose database (https://gdac.broadinstitute.org/).

### Conflict of Interest

There is no conflict of interest in with regarding to authors’ contributions

### Funding

The project was completed by using first author’s own funds. No external funding was involved.

### Author’s Contributions

The project was implemented and paper was written by first author. Second author provided guidance for forming the workflow and methodology of the study.

## Acknowledgements

This study could not have been without the guidance and support of my supervisor Dr. Saira Karim.

## Notes

https://gdac.broadinstitute.org/

### Competing Interest Statement

The authors have declared no competing interest.

